# CPLfold: Chimeric and Pseudoknot-capable almost Linear-time RNA Secondary Structure Prediction

**DOI:** 10.64898/2026.02.12.704779

**Authors:** Ke Wang, Grzegorz Kudla, Shay B. Cohen

## Abstract

Motivation RNA structure plays a central role in how transcripts function, but inferring it reliably remains difficult, especially when pseudoknots need to be part of the prediction. Chemical probing experiments provide additional signals, yet these signals do not directly identify base pairing partners. RNA proximity ligation provides direct evidence of base pairing, but balancing this evidence with pseudoknot prediction accuracy and scalability of structure prediction for long sequences remains challenging.

Results We present CPLfold, a fast and flexible RNA folding method that combines thermodynamic modeling with chimeric evidence from RNA cross-linking and ligation experiments, while naturally supporting pseudoknots. CPLfold scales to long sequences and recovers more accurate global structures and long-range interactions than existing approaches across multiple benchmarks such as COMRADES and IRIS. By tuning two simple trade-off parameters (*α, β*) the method allows flexibility in the level of incorporating chimeric evidence and asserting pseudoknots.

Availability and Implementation Source code and scripts are available at https://github.com/Vicky-0256/CPLfold.

## 1. Introduction

RNA secondary structure determines, among other things, transcript function, influencing splicing, translation, and sub-cellular localization (1). Developing accurate and scalable methods for RNA structure inference is important for explaining transcriptomic regulatory mechanisms (2), understanding viral genome organization (3, 4), and enabling RNA-targeted therapeutic interventions (5).

For *in vivo* structure determination, chemical probes such as SHAPE and icSHAPE have been widely used to guide folding, as they provide nucleotide-resolution reactivity readouts (6, 7). Many methods have incorporated reactivity signals into thermodynamic folding through pseudo-energy terms, soft constraints, or probabilistic models, thereby improving consistency with experimental data across many datasets (8, 9). However, these signals inherently reflect *local accessibility and pairing propensity* rather than *pairwise* information specifying which nucleotides pair with which. The same reactivity pattern can correspond to multiple structurally distinct but nearly equivalent interpretations, leading to substantial structural ambiguity, especially with long-range interactions or conformational heterogeneity (6, 10). Furthermore, chemical reactivity is subject to confounding factors including protein occlusion, RNA modifications, and experimental biases; when reactivity systematically deviates from the true pairing state, reliance on SHAPE-like evidence alone often fails to reliably recover critical long-range base pairs and complex topologies.

In recent years, cross-linking and ligation assays have provided observations more directly related to the structural variables of interest. Methods such as CLASH (11), PARIS (12) and COMRADES (3) generate chimeric reads through cross-linking and proximity ligation, thereby reporting RNA–RNA spatial proximities at transcriptome scale and revealing long-range base-pairing partners. Compared to the local reactivity profiles of SHAPE, these data are in principle better suited to pinpoint long-range interactions, cross-domain pairings, and distal structural organization within viral genomes. Nevertheless, cross-linking and ligation data suffer from similar confounding factors as chemical probing, and in addition, signals based on ensembles of different structures may appear to be mutually inconsistent.

Beyond evidence integration, computational complexity further limits the use of cross-linking and ligation evidence for long RNAs. Pseudoknots, formed when nucleotides within a hairpin loop base-pair with a distal single-stranded region to create interleaved stems (13), and other non-nested topologies are prevalent and functionally important in real-world RNA structures, yet permitting pseudoknots significantly increases computational complexity, rendering many classical algorithms impractical for long transcripts and infeasible for genome-scale RNAs. Existing frameworks related to cross-linking/ligation and chemical probing have made progress along different dimensions but face clear limitations. For instance, IRIS (10) relates PARIS/icSHAPE data to structural ensembles and provides a probabilistic modeling approach, but relies on pseudoknot-free cubic-time algorithms and restricts analysis to sequences under 500 nucleotides. The COMRADES pipeline (3) integrates chimeric evidence through constraint-driven thermodynamic folding, but its hard constraint mechanism is sensitive to noise and does not explicitly optimize for pseudoknotted topologies. In practice, one must often sacrifice at least one of the three: **scalability** (long sequences and large-scale data), **structural expressivity** (pseudoknot-capable outputs), and **robust experimental integration** (stable performance under conflicting evidence).

To address these challenges, we introduce **CPLfold** (**C**himeric and **P**seudoknot-capable almost **L**inear-time **fold**): a scalable folding framework built upon the LinearFold (14) algorithm and enables pseudoknot prediction with a near-linear beam-search decoding core (under bounded beam size and candidate limits) while integrating chimeric evidence from cross-linking and ligation assays with thermodynamic energy through soft constraints. CPLfold converts chimeric support into pairwise support scores and couples them with thermodynamic energy through an explicit trade-off parameter *α*, enabling the model to balance experimental consistency and physical stability in a controlled manner even when evidence is sparse and conflicting. At the algorithmic level, we introduce a two-phase cascade on top of LinearFold’s beam-search decoding, decoupling scaffold candidate generation from crossing-pair extension, followed by global re-ranking of candidate structures, thereby achieving a favorable balance between scalability and structural expressivity. Importantly, *α* is designed as an interpretable control knob that supports objective-driven tuning for different application goals (e.g., probing consistency or chimeric consistency metrics), rather than enforcing a single universal setting. We further introduce an optional selection-bias parameter *β* that controls how readily pseudoknot augmentations are retained during final selection, enabling a tunable trade-off between conservative pseudoknot calling and increased pseudoknot sensitivity.

We systematically validate CPLfold in three settings: (i) on a large curated pseudoknot benchmark (ArchiveII) to assess accuracy, coverage, and scalability; (ii) on the PARIS/icSHAPE IRIS benchmark using a strict leave-one-out protocol with icSHAPE likelihood-based *α* selection; and (iii) in a genomescale Zika virus case study demonstrating long-sequence folding capability and objective-driven *α* tuning based on chimeric support to improve long-range interaction consistency.

## 2. Methods

To enable scalable RNA secondary structure prediction guided by crosslinking–ligation evidence, we extend LinearFold (14) with a two-phase cascade and a soft evidence integration scheme. We first generate a small set of diverse nested scaffolds, then refine each scaffold under pairing constraints to introduce crossing pairs, and finally select the output from a bounded candidate pool using a pseudoknot-capable energy evaluation. Candidate generation in Phase 1/2 minimizes the evidence-augmented nested objective *E*_aug_ using a nested-energy backend (ViennaRNA or CONTRAfold), whereas final selection re-ranks fixed candidates with a pseudoknot-capable energy model *E*_PK_ (DP09 by default; other parameterizations supported).

**CPLfold** (**C**himeric and **P**seudoknot-capable **L**inear-time

### Algorithm 1.

Pseudocode for the CPLfold Algorithm

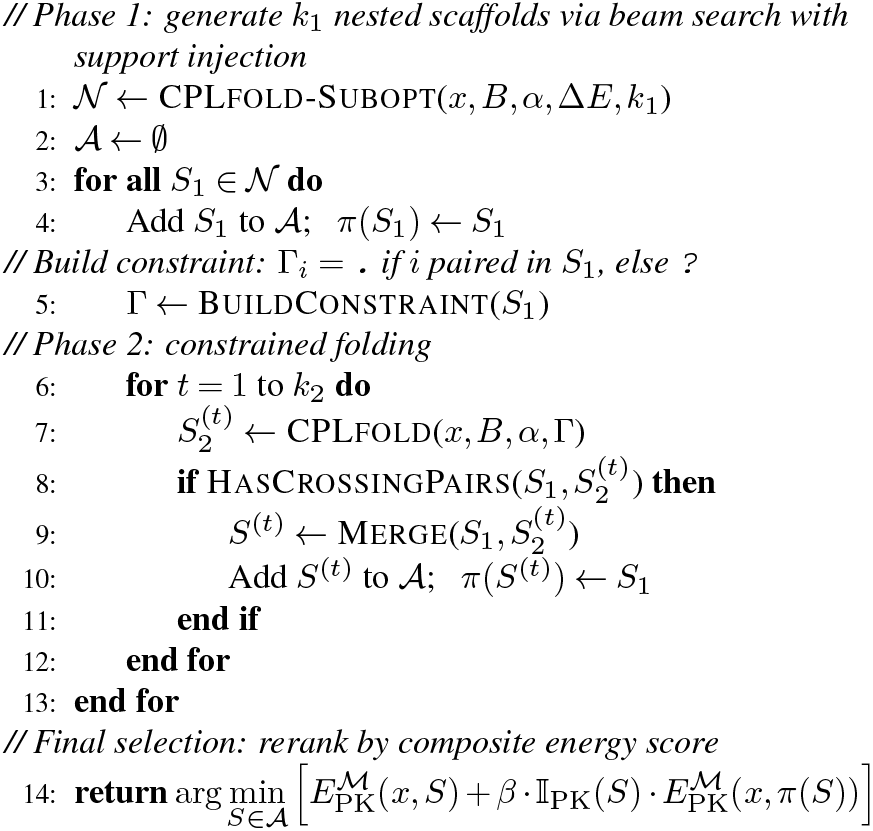

fold) takes as input an RNA sequence *x* = *x*_1_ … *x*_*n*_ and a pairwise support matrix *B* derived from proximity ligation experiments (e.g., PARIS(12), COMRADES(3)), and outputs a secondary structure prediction *S* that permits single-layer crossing extensions (nested pairs denoted by (), crossing pairs by an additional layer []), along with the final selection score under a pseudoknot-capable energy model (DP09(15) by default).

We unify thermodynamics and chimeric evidence through an additive augmented energy, with a single parameter *α* controlling the trade-off strength, thereby rewarding supported pairs without relying on brittle hard constraints.

During candidate generation in both phases, CPLfold minimizes the augmented energy

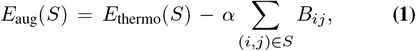

where *E*_thermo_(*S*) is the thermodynamic score, *B*_*ij*_ ≥ 0 denotes the chimeric evidence support at position (*i, j*), and *α* ≥ 0 is the evidence weight. When *α* = 0, the method reduces to LinearFold-style decoding without experimental information; when *α >* 0, it favors retaining pairs with chimeric support. Construction of the support matrix *B* is described in Section E.

### A. Phase 1: Nested folding and *k*-best enumeration

Phase 1 builds upon LinearFold’s left-to-right beam-search decoding (14): we leave the dynamic programming states, transition rules, and pruning strategies unchanged, replacing only the scoring objective from pure thermodynamic energy to the augmented energy *E*_aug_ (Eq. 1), thereby allowing chimeric evidence to influence the search trajectory as a soft reward. The output of this phase is not a single structure but a set of high-quality nested candidates within an energy band, which serve as scaffolds for constrained refinement in Phase 2.

Only Watson–Crick (A–U, G–C) and wobble (G–U) pairs are allowed, and hairpin loops must satisfy *j* − *i*> 3. CPLfold maintains a beam of at most *w* partial states at each position. When a state transition creates a new base pair (*i, j*), we subtract *αB*_*ij*_ from the local energy, ensuring that each pair’s support score is counted exactly once.

We support two nested-energy backends: ViennaRNA Turner 2004 parameters (16) and CONTRAfold (17); support scores are injected in the same manner for both.

To limit search cost while reducing dependence on any single scaffold, we run a lightweight inside–outside pass in the same beam-pruned space and apply a Zuker-style band to trace up to *k*_1_ distinct scaffolds, then remove near-duplicates using a length-adaptive window.

After decoding, we run Inside–Outside on the same beam-pruned space to enumerate suboptimal scaffolds with energies close to the minimum augmented energy (MFE = *E*_aug_(*S*^∗^)). Let 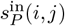 and 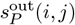 denote the inside/outside augmented energies for the paired state (*i, j*); we retain anchors satisfying the Zuker(18) band criterion:

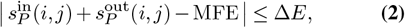

We then sort anchors by 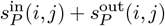 in ascending order and trace back from each anchor to generate a complete nested scaffold. To suppress near-duplicates, we apply length-adaptive window deduplication to anchors, ultimately retaining at most *k*_1_ scaffolds for Phase 2 refinement.

### B. Phase 2: Constrained folding

To add crossing pairs without disrupting the nested scaffold, we rerun the same decoder under a constraint that forbids scaffold-paired positions from forming new base pairs.

Our two-phase strategy is inspired by the hierarchical folding hypothesis: nested secondary structures often form stable scaffolds first, with additional long-range interactions (including pseudoknots) arising subsequently without substantially disrupting the scaffold (19, 20).

Given a first-phase scaffold *S*_1_, we construct a constraint string Γ ∈ {., ?}^*n*^:

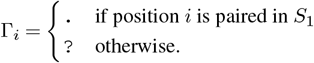

where. indicates that position *i* is forbidden from pairing in Phase 2, and ? allows pairing or remaining unpaired. Phase 2 runs the same beam-search decoder as Phase 1 under constraint Γ: if a transition would introduce base pair (*i, j*) and Γ_*i*_ =. or Γ_*j*_ =., that transition is skipped. Phase 2 uses the **same** support matrix *B* and weight *α* as Phase 1 to ensure consistent evidence guidance across both phases.

In this work, we set *k*_2_ = 1, meaning each scaffold undergoes only one constrained refinement; increasing *k*_2_ can serve as an extension to expand the search space (e.g., enumerating Phase 2 suboptimal structures or progressively relaxing a few constraints), which we do not systematically explore here.

### C. Merging and pseudoknot detection

To construct pseudoknot candidates from the two phases, we detect crossing pairs between scaffold and refinement outputs and merge them into a single structure representation.

Given the scaffold pair set *S*_1_ and the second-phase pair set *S*_2_, if there exist (*i, j*) ∈ *S*_1_ and (*k, ℓ*) ∈ *S*_2_ satisfying

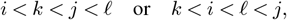

we determine that a pseudoknot (crossing pairs) has been formed.

When crossing pairs are detected, we retain first-phase pairs as () and add second-phase pairs as [], provided they do not conflict with the scaffold (i.e., both endpoints are unpaired in *S*_1_), thereby forming the merged structure *S*. We use only a single additional bracket layer to represent one crossing extension atop the nested scaffold, consistent with the target topology range of the two-phase cascade.

### D. Candidate scoring and final selection

To avoid systematically introducing spurious pseudoknots, CPLfold decouples *candidate proposal* from *final selection*. The twophase cascade may produce crossing-pair augmentations, but it is not forced to output a pseudoknotted structure: for every scaffold *S*_1_, the nested scaffold itself is always retained as a fallback candidate, and whether an augmented pseudoknot is selected is determined by thermodynamic plausibility.

For each scaffold *S*_1_, CPLfold adds it to the candidate pool 𝒜 ; if a merged pseudoknotted structure *S* is produced, it is also added to 𝒜. Thus | 𝒜 | ≤ *k*_1_(1 + *k*_2_); with the default *k*_2_ = 1, we have |𝒜| ≤ 2*k*_1_.

Since pseudoknotted structures cannot be scored by nested decompositions, we evaluate each fixed candidate using a pseudoknot-capable energy model, denoted *E*_PK_(*x, S*); we use the same *E*_PK_ parameterization within a run for scoring both candidates and their associated scaffolds. We consider three widely adopted energy parameterizations: DP09 (Dirks–Pierce)(15), CC09 (Cao–Chen)(21), and RE(13). The DP model extends standard nearest-neighbor free energy parameters with pseudoknot-specific penalty terms; the CC model provides a more detailed treatment of H-type pseudoknot thermodynamics through refined loop entropy and coaxial stacking contributions, reverting to DP parameters where CC-specific values are undefined (22).

We further introduce a scalar hyperparameter *β* ≥ 0 to optionally bias selection toward candidates with crossing-pair augmentations. Let 𝕀_PK_(*S*) ∈ { 0, 1 } indicate whether *S* contains crossing pairs, and let *π*(*S*) denote the phase-1 scaffold associated with *S* (with *π*(*S*_1_) = *S*_1_ for the scaffold itself). We define the composite selection score

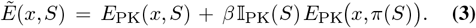

Because free energies are typically negative, increasing *β* effectively rewards pseudoknotted augmentations built on thermodynamically stable scaffolds, making such candidates more likely to be selected.

The final output is selected by directly choosing the minimum-score candidate from the pool, without additional tructure search:

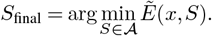

Unless otherwise noted, we set *β* = 0, which reduces the selection rule to pure *E*_PK_(*x, S*) evaluation.

### E. Chimeric data integration

To transform sparse chimeric reads into a model-compatible signal, we aggregate evidence into a pairwise support matrix and optionally apply a log transform to compress dynamic range, then use it as the soft support term in Eq. 1.

We first map chimeric reads to the target transcript using the program hyb (23), representing each read as two arms (left arm [*ll, lr*) and right arm [*rl, rr*)). For each read, we perform cross-arm duplex inference and extract canonical base pairs, accumulating them into a symmetric count matrix *C* (*C*_*ij*_ = *C*_*ji*_). The support matrix *B* is then derived from *C* as *B* = *C* (for raw counts) or *B* = log (1 + *C*) for log-compressed, with log applied coordinate-wise on *C*. Log compression is suitable for *α* selection across multiple sequences to balance dynamic range; raw counts are more appropriate for objective optimization on single long sequences. To maintain scalability, *B* is stored as a sparse mapping and queried only when beam search examines specific pairs.

### F. Computational complexity

To quantify scalability, we analyze time complexity in terms of sequence length *n*, beam size *w*, and candidate limits (*k*_1_, *k*_2_), showing that the decoding passes scale near-linearly in *n* under bounded beam size and candidate limits, while the remaining steps are linear in the bounded candidate pool size.

In a single run, CPLfold executes one first-phase decoding pass and at most *k*_1_*k*_2_ constrained second-phase decoding passes with total decoding complexity *O* ((1 + *k*_1_ *k*_2_) *n* · *w* log *w*). Phase 1 adds an additional inside–outside pass in the same beam-pruned space, which does not change the near-linear scaling under bounded *w*.

The candidate pool size satisfies | 𝒜 | ≤ *k*_1_(1 + *k*_2_); thus the total energy evaluation cost is *O* (|𝒜| · |*S* |), where | *S* | denotes the number of base pairs in *S*, which for sparse secondary structures can be approximated as *O* (|𝒜| · *n*). With the default *k*_2_ = 1, fixed *w*, and bounded candidate limits, the end-to-end pipeline maintains near-linear scaling with respect to *n*.

### G. Implementation and reproducibility

CPLfold is implemented in Python, with performance-critical kernels JIT-compiled using Numba (LLVM). Cross-arm duplex inference uses ViennaRNA (24); pseudoknot energy evaluation uses DP09 by default (CC09/RE also supported). Because JIT compilation incurs one-time startup overhead, all runtimes reported in this work are steady-state performance after warm-up.

## 3. Results

This section organizes experiments around three central claims of CPLfold. First, **Scalability:** CPLfold supports stable pseudoknot-capable folding at scale without introducing super-cubic time complexity, making it applicable to large collections of sequences. Second, **Evidence integration:** chimeric and crosslinking evidence can be incorporated into thermodynamic decoding as soft constraints through a single trade-off parameter *α*. Even in the absence of groundtruth structures, *α* can be selected in a stable manner using a held-out objective, yielding gains that generalize across sequences. Third, **Genome-scale feasibility and objective-driven tuning:** CPLfold remains computationally practical for ultra-long RNAs on the order of 10 kb, and *α* can be chosen directly by optimizing an evidence-consistency objective, leading to a substantial increase in agreement with *in vivo* evidence with only seconds of runtime overhead.

Throughout, candidate generation uses a nested-energy back-end while final selection uses a pseudoknot-capable model *E*_PK_ (DP09 by default); we choose task-native external baselines per setting and include *α* = 0 internal controls. In the *β*-biased selection rule, the scaffold term is evaluated under the same *E*_PK_ model used for final re-ranking.

Following standard practice in RNA secondary structure prediction, we evaluated structural accuracy using precision (PPV; positive predictive value), recall (SEN; sensitivity), and F1. Concretely, we extract the set of aligned base pairs from both the predicted and the reference structures, compute precision and recall on these paired sets, and report macro-averaged statistics by averaging per-sequence precision, recall, and F1. All structure metrics are computed on each method’s **successful subset**, that is, only sequences for which the method returns an output structure are scored. We additionally report **coverage** (the number of successful sequences divided by the total number of sequences) to characterize failure behavior on long RNAs. For pseudoknot detection, the positive label is defined by the presence of at least one crossing pair in the reference annotation; metrics are computed on each method’s successful subset, with coverage reported separately.

Under the IRIS setting, we adopt two unsupervised criteria from IRIS (10). **icSHAPE log-likelihood (LL):** we use the IRIS generative model to quantify the agreement between the predicted structure set and icSHAPE measurements, where larger values indicate better consistency (6, 10). Because icSHAPE measures local accessibility rather than pairwise proximity, it is orthogonal to PARIS evidence and can therefore be used to select *α* without relying on reference structures. **KL divergence (KL):** we compare the predicted base-pairing probability matrix to the normalized mutual information matrix derived by R-scape (25) (corresponding to IRIS, where smaller values are better), thereby measuring agreement with evolutionary covariation signals.

In the ZIKV setting, where a genome-wide ground-truth structure is unavailable, we evaluate predictions using a COMRADES-based consistency metric. For every pair of positions (*i, j*) along the viral genome, we use the COMRADES score *C*_*ij*_, defined as the number of chimeric reads that, when analyzed with `hybrid-min` under default settings, indicate base pairing between positions *i* and *j*(3). Let *S* denote the set of base pairs in a predicted structure. We define the chimeric consistency score as

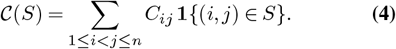

Larger 𝒞 (*S*) indicates stronger consistency with *in vivo* proximity-ligation evidence captured by COMRADES.

### A. Scalable pseudoknot-capable folding on ArchiveII

To assess whether CPLfold can stably produce pseudoknot-capable structures at scale without incurring super-cubic complexity, we conducted a sequence-only evaluation on the ArchiveII benchmark (26). We compare runtime scaling, secondary-structure accuracy, and pseudoknot (PK) detection performance, and we additionally report coverage to reflect failure cases on long sequences. ArchiveII comprises 3,975 sequences from 10 RNA families with gold-standard annotations that include pseudoknots (length 28–2,968 nt). All methods perform *de novo* prediction from sequence alone. For CPLfold, we set *α* = 0 to disable the support term so that both phase-1 and phase-2 candidate generation are driven solely by the thermodynamic backend. Unless otherwise specified, we use beam size *w*=100, Δ*E*=5.0 kcal/mol, and *k*_1_=10, and we perform one constrained refinement per scaffold (*k*_2_=1). We compare against HotKnots v2.0 (22)and RNAstructure-ProbKnot(27), and we evaluate CPLfold using ViennaRNA or CONTRAfold as the candidate-generation backend. For final selection, we rescore the fixed candidate set under three pseudoknot-capable energy parameterizations (DP09, CC09, and RE), with DP09 as the default.

#### A.1. Runtime scaling and coverage

To characterize empirical complexity, we model runtime using a power-law relationship *t* = *a* · *n*^*b*^, where *t* is runtime in seconds, *n* is sequence length, and *b* is the complexity exponent. To obtain robust estimates insensitive to outliers, we employ a binned median fitting approach: sequences are grouped into length intervals, and the median runtime within each bin is computed. Linear regression is then performed on log-transformed bin centers and median runtimes. All three methods achieve *R*^2^ *>* 0.98, indicating excellent model fit (Fig. 1).

**Fig. 1.**
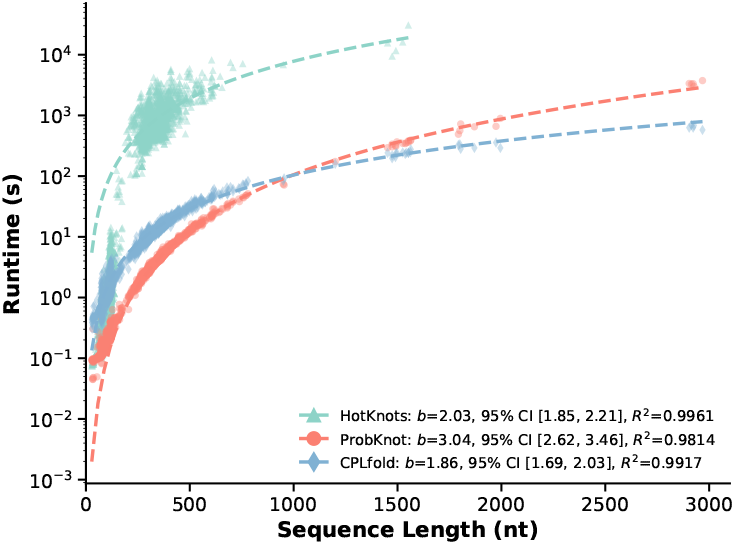
Runtime scaling on ArchiveII. Each point shows the single-sequence runtime as a function of sequence length.

HotKnots is competitive on short sequences, but becomes impractical as length increases: runtimes rise from seconds for sequences shorter than 200 nt to hundreds or thousands of seconds for 300–500 nt, and reach hour-level beyond 700 nt. HotKnots also fails to return outputs for 60 long sequences, yielding a coverage of 3,915/3,975 (98.5%). RNAstructure-ProbKnot achieves full coverage up to 2,968 nt (100%), but exhibits cubic scaling (*O*(*n*^3.04^)) and requires tens to hundreds of seconds in the 1,000–3,000 nt range. In contrast, CPLfold shows an empirical runtime scaling exponent of *b* = 1.86 on ArchiveII (Fig. 1), indicating sub-quadratic growth in practice.

#### A.2. Structure prediction accuracy

Structural accuracy is summarized in Table 1 (left). Under the same sequence-only setting, the two-phase scaffold-based generation followed by cross refinement yields consistent gains over the pseudoknot-free baseline (single-phase LinearFold). Specifically, CPLfold-Vienna (DP09) improves F1 from 0.561 to 0.606 (+8.0%), and CPLfold-CONTRAfold (DP09) improves F1 from 0.601 to 0.626 (+4.2%). On their respective successful subsets, CPLfold-CONTRAfold (DP09) also out-performs HotKnots (F1 0.626 vs. 0.562). Varying the pseudoknot energy parameterization used for *final candidate selection* produces small but consistent differences (with CC09 achieving the highest F1), suggesting that pseudoknot-aware rescoring has a controlled yet substantive effect on end-to-end quality.

**Table 1.** ArchiveII: Structure prediction and pseudoknot detection performance.

| Method | Structure |  |  | Pseudoknot |  |  |
| --- | --- | --- | --- | --- | --- | --- |
|  | PPV | SEN | F1 | PPV | SEN | F1 |
| LinearFold-Vienna | 0.537 | 0.592 | 0.561 | n/a |  |  |
| LinearFold-CONTRAFold | **0.621** | 0.596 | 0.601 |  |  |  |
| HotKnots v2.0 | 0.544 | 0.585 | 0.562 | 0.385 | 0.038 | 0.070 |
| RNAstructure-ProbKnot | 0.554 | 0.601 | 0.573 | 0.514 | 0.498 | 0.506 |
| CPLfold-Vienna (DP09) | 0.582 | 0.639 | 0.606 | 0.147 | 0.015 | 0.027 |
| CPLfold-CONTRAFold (DP09) | 0.618 | 0.643 | 0.626 | 0.581 | 0.099 | 0.170 |
| CPLfold-CONTRAFold (DP09, $\beta=0.46$) | 0.593 | 0.622 | 0.604 | **0.666** | **0.817** | **0.734** |
| CPLfold-CONTRAFold (CC09) | 0.618 | **0.643** | **0.626** | 0.565 | 0.097 | 0.165 |
| CPLfold-CONTRAFold (CC09, $\beta=0.46$) | 0.590 | 0.619 | 0.600 | 0.653 | 0.804 | 0.721 |
| CPLfold-CONTRAFold (RE) | 0.586 | 0.626 | 0.602 | 0.529 | 0.060 | 0.108 |
| CPLfold-CONTRAFold (RE, $\beta=0.62$) | 0.559 | 0.600 | 0.575 | 0.615 | 0.818 | 0.702 |

#### A.3. Pseudoknot detection performance

Pseudoknot detection results are reported in Table 1 (right). Under the default setting (*β* = 0), CPLfold-CONTRAfold (DP09) exhibits a conservative “high-precision, low-recall” pattern relative to RNAstructure-ProbKnot, achieving precision 0.581 (vs. ProbKnot: 0.514) but recall of only 0.099 (vs. ProbKnot: 0.498), indicating that under purely energy-driven final selection, the model retains only high-confidence pseudoknots. Adjusting the selection bias parameter *β* (Eq. 3) substantially improves pseudoknot detection. As shown in Fig. 2, under all three parameterizations (DP09, CC09, RE), pseudoknot detection F1 increases rapidly with *β* and saturates around *β* ≈ 0.2–0.4: DP09 reaches a peak pseudoknot F1 of 0.734 at *β* = 0.46, CC09 achieves 0.721 at *β* = 0.46, and RE achieves 0.702 at *β* = 0.62. Meanwhile, overall structure F1 remains largely stable, with only a slight decrease (e.g., DP09 drops from 0.626 to approximately 0.59). Thus, *β* provides a controllable trade-off: a modest sacrifice in overall structural accuracy yields a substantial gain in pseudoknot detection. The default *β* = 0 is appropriate when overall structure accuracy is prioritized, whereas grid search over *β* can identify the optimal value for tasks emphasizing pseudoknot sensitivity.

**Fig. 2.**
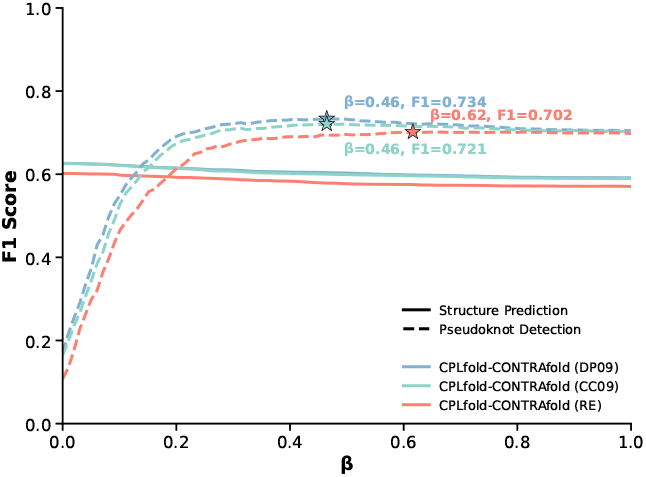
Effect of *β* on F1 on ArchiveII.

### B. Chimera-guided folding on the IRIS benchmark

To test whether the proposed soft integration yields generalizable gains under real crosslinking and ligation evidence, we reproduced the IRIS benchmark workflow on the 11-transcript dataset and applied a strict leave-one-out protocol. We select *α* using held-out icSHAPE log-likelihood, then evaluate KL, LL, and F1, and compare against IRIS and the COMRADES pipeline. We follow IRIS (10) for dataset construction, preprocessing, and evaluation definitions, using the same set of 11 RNAs that have both PARIS and icSHAPE signals (length 79–330 nt). To compress the dynamic range of read counts and balance contributions across transcripts of different lengths, we apply a log-compressed support transform, *B*_*ij*_ = log(1 + *C*_*ij*_). In this experiment, CPLfold uses CONTRAfold as the phase-1 and phase-2 candidate-generation backend and DP09 for the final energy-based candidate evaluation.

#### B.1. A strict leave-one-out protocol for selecting α using held-out LL

Because the true *in vivo* structure ensembles are unknown, we select *α* without using curated reference structures and adopt a strict leave-one-out scheme to prevent information leakage. For each held-out RNA *t*, we search over *α* on the remaining 10 RNAs and optimize the sum of training-fold icSHAPE log-likelihoods, obtaining 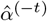. We then fix 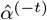 and apply it to the held-out RNA *t* to compute test metrics. Formally,

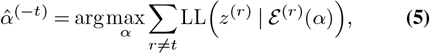

where *z*^(*r*)^ denotes the icSHAPE score vector of RNA *r*, and ℰ^(*r*)^(*α*) denotes the predicted structure set under *α*, which corresponds to the top-ranked candidate from CPLfold. This protocol is used only to select the global trade-off parameter *α* and does not involve training model parameters.

#### B.2. Effect of α under grid evaluation

To examine how evidence weighting influences different evaluation axes, we swept *α* over a grid and evaluated CPLfold on each of the 11 RNAs (Fig. 3). The resulting trajectories reveal substantial heterogeneity across transcripts and across metrics. In particular, the *α* values that improve structural F1 against curated references are often not the ones that maximize icSHAPE log-likelihood, and the KL divergence to covariation signals can follow yet another pattern. Thus, the three criteria are not aligned and no single *α* is uniformly optimal across sequences, motivating the use of *α* as a tunable control knob chosen according to the task objective.

**Fig. 3.**
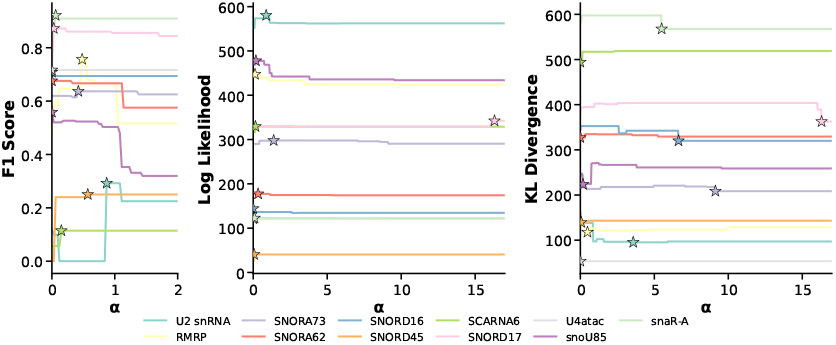
Effect of the trade-off parameter *α* on CPLfold across the 11 IRIS PARIS transcripts. Each colored line corresponds to one RNA and reports performance as *α* varies.

#### B.3. Overall performance under strict leave-one-out selection

Across the 11 leave-one-out folds, the selected 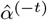 values are highly concentrated. Specifically, 8 of 11 transcripts select *α* ≈ 0.5131 (median 0.5131; range 0.3338– 0.5655). Under the strict leave-one-out selection of *α*, CPLfold achieves higher structural accuracy (Table 2). Among all compared methods, CPLfold-3 attains the highest mean F1 of 0.560, exceeding IRIS-1 (0.516), IRIS-2 (0.446), and IRIS-3 (0.415). At the same time, CPLfold-3 attains an LL of 277.6, which is comparable to IRIS-1 (276.0) and slightly below IRIS-3 (280.3)(Table 4). For KL, LinearFold yields the lowest value (243.7), while CPLfold and IRIS fall in a similar range of 253–283(Table 3).

**Table 2.** Per-RNA F1 across methods under leave-one-out *α* selection.

| RNA | Len | Reads | MFE | LF | COMRADES | IRIS |  |  | CPLfold (ours) |  |  |
| --- | --- | --- | --- | --- | --- | --- | --- | --- | --- | --- | --- |
|  |  |  |  |  |  | IRIS-1 | IRIS-2 | IRIS-3 | CPLfold-1 | CPLfold-2 | CPLfold-3 |
| U2 snRNA | 192 | 1142 | <u>0.099</u> | 0.000 | <u>0.111</u> | 0.087 | 0.045 | 0.039 | 0.000 | 0.000 | 0.093 |
| RMRP | 264 | 5919 | <u>0.490</u> | 0.434 | 0.366 | <u>0.553</u> | 0.453 | 0.357 | <u>0.756</u> | <u>0.756</u> | <u>0.756</u> |
| SNORA73 | 207 | 9592 | <u>0.619</u> | 0.540 | 0.478 | 0.491 | 0.478 | 0.398 | 0.614 | 0.617 | <u>0.623</u> |
| SNORA62 | 153 | 1038 | <u>0.676</u> | 0.716 | 0.575 | 0.568 | 0.532 | 0.485 | 0.667 | 0.658 | 0.661 |
| SNORD16 | 99 | 1964 | <u>0.694</u> | 0.698 | 0.588 | 0.714 | 0.586 | 0.561 | 0.694 | 0.660 | 0.648 |
| SNORD45 | 79 | 1641 | 0.000 | 0.250 | 0.242 | <u>0.261</u> | <u>0.255</u> | 0.249 | 0.247 | 0.247 | 0.247 |
| SCARNA6 | 275 | 1400 | 0.057 | <u>0.099</u> | 0.056 | 0.043 | 0.081 | 0.087 | <u>0.114</u> | <u>0.114</u> | <u>0.114</u> |
| SNORD17 | 237 | 1307 | 0.749 | 0.783 | 0.837 | <u>0.884</u> | 0.799 | 0.767 | <u>0.867</u> | <u>0.867</u> | <u>0.867</u> |
| U4atac | 127 | 2635 | 0.716 | 0.393 | 0.627 | 0.716 | 0.396 | 0.394 | 0.716 | <u>0.722</u> | <u>0.716</u> |
| snoU85 | 330 | 1257 | <u>0.547</u> | 0.470 | 0.398 | <u>0.540</u> | 0.519 | 0.514 | 0.523 | 0.523 | 0.523 |
| snaR-A | 115 | 1954 | 0.842 | <u>0.849</u> | 0.513 | 0.816 | 0.773 | 0.715 | <u>0.909</u> | <u>0.909</u> | <u>0.909</u> |
| **Average** | — | — | 0.499 | 0.449 | 0.436 | 0.516 | 0.446 | 0.415 | <u>0.556</u> | 0.552 | <u>0.560</u> |

For each RNA $t$ , $\hat{\alpha}^{(-t)}$ is selected on the remaining 10 RNAs by maximizing icSHAPE log-likelihood (LL) and then applied to the held-out RNA $t$ for evaluation. The suffix $k$ in CPLfold- $k$ (resp. IRIS- $k$ ) denotes the number of top-ranked candidates used as the structure ensemble for evaluation. LF denotes LinearFold-CONTRAFold; MFE results are consistent with LinearFold-Vienna. Tables 3 and 4 use the same configuration and report KL and LL, respectively. Best two values per row are underlined.

**Table 3.**
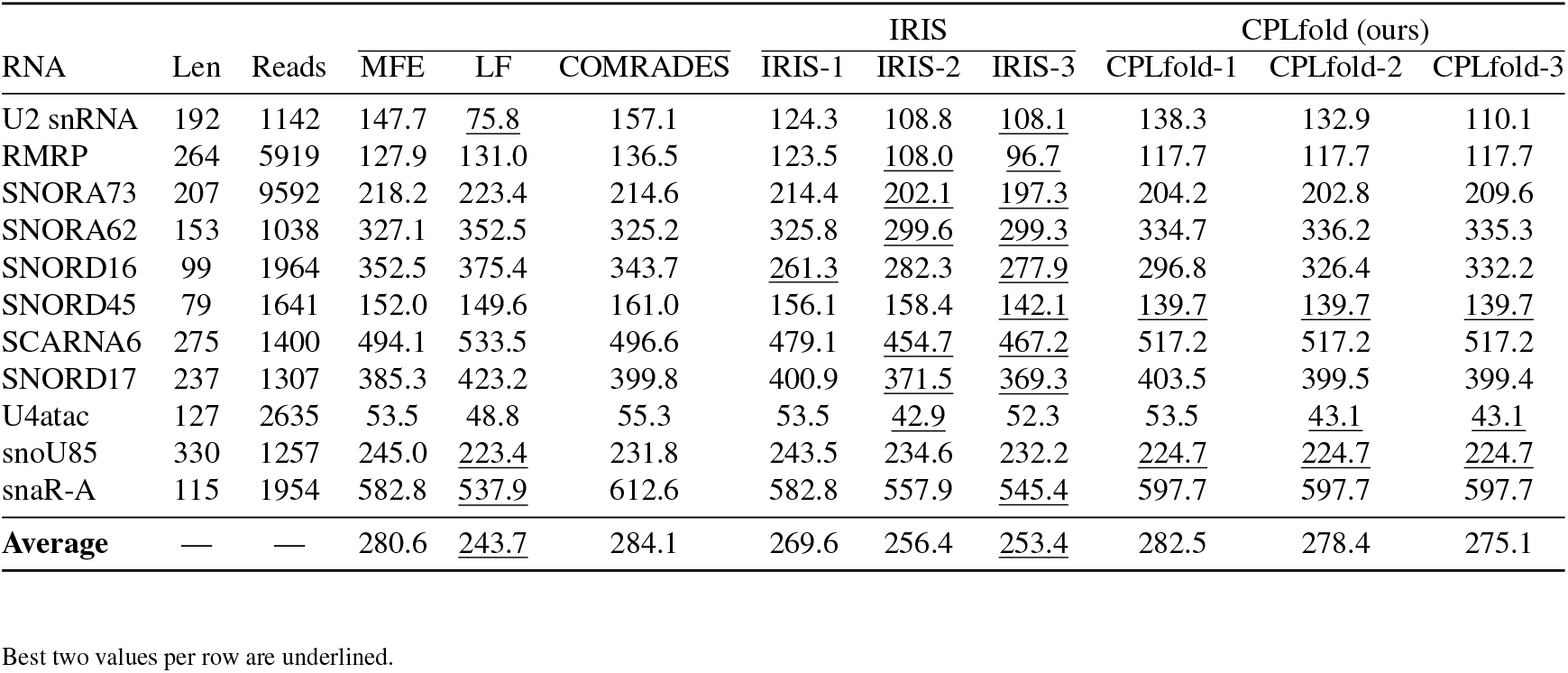
Per-RNA KL divergence across methods under leave-one-out *α* selection.

| RNA | Len | Reads | MFE | LF | COMRADES | IRIS |  |  | CPLfold (ours) |  |  |
| --- | --- | --- | --- | --- | --- | --- | --- | --- | --- | --- | --- |
|  |  |  |  |  |  | IRIS-1 | IRIS-2 | IRIS-3 | CPLfold-1 | CPLfold-2 | CPLfold-3 |
| U2 snRNA | 192 | 1142 | 147.7 | <u>75.8</u> | 157.1 | 124.3 | 108.8 | <u>108.1</u> | 138.3 | 132.9 | 110.1 |
| RMRP | 264 | 5919 | 127.9 | 131.0 | 136.5 | 123.5 | <u>108.0</u> | <u>96.7</u> | 117.7 | 117.7 | 117.7 |
| SNORA73 | 207 | 9592 | 218.2 | 223.4 | 214.6 | 214.4 | <u>202.1</u> | <u>197.3</u> | 204.2 | 202.8 | 209.6 |
| SNORA62 | 153 | 1038 | 327.1 | 352.5 | 325.2 | 325.8 | <u>299.6</u> | <u>299.3</u> | 334.7 | 336.2 | 335.3 |
| SNORD16 | 99 | 1964 | 352.5 | 375.4 | 343.7 | <u>261.3</u> | 282.3 | <u>277.9</u> | 296.8 | 326.4 | 332.2 |
| SNORD45 | 79 | 1641 | 152.0 | 149.6 | 161.0 | 156.1 | 158.4 | <u>142.1</u> | <u>139.7</u> | <u>139.7</u> | <u>139.7</u> |
| SCARNA6 | 275 | 1400 | 494.1 | 533.5 | 496.6 | 479.1 | <u>454.7</u> | <u>467.2</u> | 517.2 | 517.2 | 517.2 |
| SNORD17 | 237 | 1307 | 385.3 | 423.2 | 399.8 | 400.9 | <u>371.5</u> | <u>369.3</u> | 403.5 | 399.5 | 399.4 |
| U4atac | 127 | 2635 | 53.5 | 48.8 | 55.3 | 53.5 | <u>42.9</u> | 52.3 | 53.5 | <u>43.1</u> | <u>43.1</u> |
| snoU85 | 330 | 1257 | 245.0 | <u>223.4</u> | 231.8 | 243.5 | 234.6 | 232.2 | <u>224.7</u> | <u>224.7</u> | <u>224.7</u> |
| snaR-A | 115 | 1954 | 582.8 | <u>537.9</u> | 612.6 | 582.8 | 557.9 | <u>545.4</u> | 597.7 | 597.7 | 597.7 |
| **Average** | — | — | 280.6 | <u>243.7</u> | 284.1 | 269.6 | 256.4 | <u>253.4</u> | 282.5 | 278.4 | 275.1 |

Best two values per row are underlined.

**Table 4.** Per-RNA icSHAPE log-likelihood LL across methods under leave-one-out *α* selection.

| RNA | Len | Reads | MFE | LF | COMRADES | IRIS |  |  | CPLfold (ours) |  |  |
| --- | --- | --- | --- | --- | --- | --- | --- | --- | --- | --- | --- |
|  |  |  |  |  |  | IRIS-1 | IRIS-2 | IRIS-3 | CPLfold-1 | CPLfold-2 | CPLfold-3 |
| U2 snRNA | 192 | 1142 | 551.5 | 573.8 | 544.0 | 579.5 | <u>581.8</u> | <u>581.5</u> | 573.5 | 572.7 | 579.7 |
| RMRP | 264 | 5919 | 435.4 | 402.3 | 430.1 | 444.3 | <u>459.7</u> | <u>459.2</u> | 440.5 | 440.5 | 440.5 |
| SNORA73 | 207 | 9592 | 298.2 | 296.8 | 292.0 | 298.3 | 295.4 | 294.2 | 297.7 | 297.7 | 297.7 |
| SNORA62 | 153 | 1038 | 176.9 | 177.2 | <u>185.6</u> | 176.9 | 173.3 | 172.7 | 177.3 | 177.3 | <u>177.4</u> |
| SNORD16 | 99 | 1964 | <u>144.8</u> | 133.6 | 135.8 | <u>144.8</u> | 141.4 | <u>143.1</u> | 136.4 | 136.4 | 136.4 |
| SNORD45 | 79 | 1641 | 25.4 | 23.0 | 17.9 | <u>40.6</u> | <u>40.6</u> | <u>48.7</u> | <u>40.6</u> | <u>40.6</u> | <u>40.6</u> |
| SCARNA6 | 275 | 1400 | 316.3 | 310.9 | 307.9 | 316.9 | 322.3 | <u>323.2</u> | <u>329.4</u> | <u>329.4</u> | <u>329.4</u> |
| SNORD17 | 237 | 1307 | 321.2 | 318.9 | 329.1 | <u>330.3</u> | 329.8 | <u>335.7</u> | <u>330.3</u> | <u>330.3</u> | <u>330.3</u> |
| U4atac | 127 | 2635 | <u>121.4</u> | 117.4 | 121.3 | <u>121.4</u> | 119.8 | <u>130.7</u> | <u>121.4</u> | <u>121.4</u> | <u>121.4</u> |
| snoU85 | 330 | 1257 | 468.7 | <u>473.8</u> | 444.7 | 467.4 | 473.7 | <u>473.8</u> | <u>477.5</u> | <u>477.5</u> | <u>477.5</u> |
| snaR-A | 115 | 1954 | 115.0 | <u>122.3</u> | 106.6 | 115.0 | 121.1 | 120.8 | <u>122.8</u> | <u>122.8</u> | <u>122.8</u> |
| **Average** | — | — | 269.7 | 274.3 | 266.0 | 276.0 | <u>278.3</u> | <u>280.3</u> | 276.3 | 277.0 | 277.6 |

Best two values per row are underlined.

### C. Genome-scale folding of the Zika virus RNA

To demonstrate genome-scale scalability with objective-driven tuning, we use the 10,807-nt Zika virus (ZIKV) genome. At this length, IRIS-style *O* (*n*^3^) folding is impractical. Using the ZIKV COMRADES dataset (3) (1.7 million non-redundant chimeric reads), we construct the chimeric count matrix *C* and tune *α* by directly maximizing the COMRADES chimeric-consistency objective 𝒞 (*S*) under the same evidence input.

We implemented the COMRADES pipeline (3) on the full-length 10,807-nt ZIKV genome (i.e., folding the complete sequence rather than segments). Briefly, chimeric-read evidence was aggregated into stems, the top 75 highest-scoring stems were retained as a constraint pool, and candidate structures were produced by repeatedly resampling this pool and performing one ViennaRNA constrained fold per resampling. Using this configuration, we generated 1,000 COMRADES candidates; under a single-CPU timing convention, one constrained fold took ∼ 5 hours (the full set was produced via parallel execution).

Fig. 4 shows that the 1,000 COMRADES candidates occupy a relatively narrow region; the best achieves 𝒞 (*S*) = 2.82 × 10^6^ at Δ*G*_DP09_ = − 2181.0 kcal/mol. In contrast, CPLfold provides a controllable evidence–energy trade-off: at *α* = 0 (thermodynamics-only), the prediction is driven purely by thermodynamic stability (Δ*G*_DP09_ = − 1957.1 kcal/mol) but achieves lower chimeric support (𝒞 (*S*) = 1.25 × 10^6^), whereas increasing *α* markedly improves 𝒞 (*S*) with modest changes in Δ*G*_DP09_. The maximum occurs at *α* = 0.0027, where CPLfold reaches 𝒞 (*S*) = 3.35 × 10^6^, exceeding the best COMRADES candidate by ∼ 19%. Thus, at genome scale and without ground truth, optimizing 𝒞 enables reproducible gains in *in vivo* evidence concordance without expensive randomized constraint sampling.

**Fig. 4.**
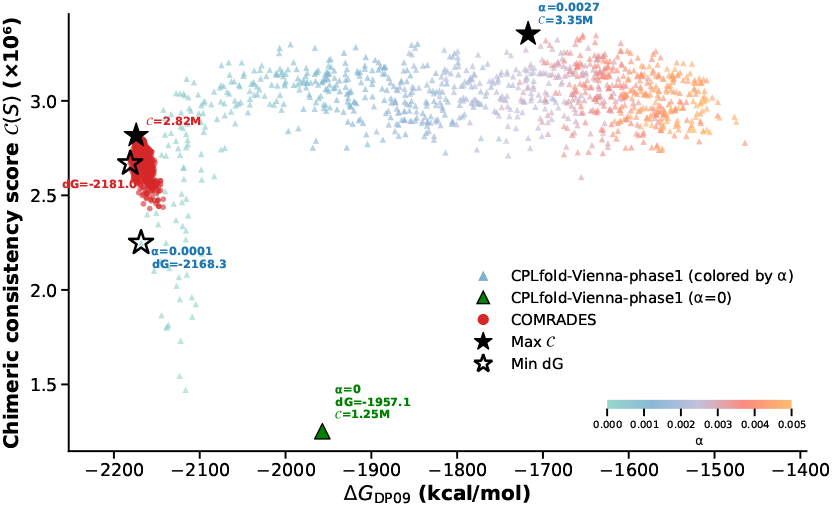
Comparison of CPLfold and COMRADES on the complete ZIKV genome (10,807 nt). 𝒞 (*S*): Chimeric Support (Eq. 4); higher values indicate stronger concordance with *in vivo* proximity-ligation evidence and serve as the primary comparison metric.

Starting from *α* = 0.0027, Phase 2 introduces crossing-pair augmentations and yields 𝒞 (*S*) = 3,363,195. The final structure has Δ*G*_DP09_ = − 946.32 kcal/mol (reported for completeness), but in this *in vivo* case study we treat evidence concordance 𝒞 (*S*) as the primary signal, since thermodynamic scores may not fully reflect cellular constraints shaped by kinetics and protein interactions(6, 28).

## 4. Discussion

We introduced CPLfold, a scalable framework for RNA secondary structure prediction that integrates crosslinking– ligation evidence and supports pseudoknot-capable decoding at practical scale. Three evaluations support three main observations. First, under sequence-only conditions, CPLfold improves structural accuracy relative to pseudoknot-free baselines while maintaining high coverage, and it exhibits sub-quadratic end-to-end scaling on ArchiveII (*b* ≈ 1.86). Second, chimeric evidence can be incorporated as soft constraints controlled by a single trade-off parameter *α*, providing an interpretable balance between thermodynamic stability and evidence concordance. Notably, *α* can be selected using held-out objectives such as icSHAPE log-likelihood (LL), enabling tuning even when ground-truth structures are unavailable. Third, in the genome-scale ZIKV case study, CPLfold remains feasible at the 10 kb scale and supports reproducible, evidence-driven tuning by directly optimizing the chimeric consistency objective 𝒞 (*S*), which improves agreement with *in vivo* COMRADES readouts.

The methodological key of CPLfold lies in transforming the unbounded search over general pseudoknots into reranking over a finite candidate set. Phase 1 enumerates a small number of diverse nested scaffolds; Phase 2 proposes crossing-pair augmentations under scaffold constraints, thereby increasing structural expressivity without expanding the search space to arbitrary pseudoknots. Global rescoring and selection over a fixed candidate pool keep both computational budget and output topology under control. The ArchiveII results confirm that this bounded-candidate design achieves robust end-to-end performance while maintaining high coverage. It should be emphasized that the reported *b* ≈ 1.86 is an empirical exponent measured on end-to-end runtime over a finite length range; it includes overhead from candidate enumeration, deduplication, and rescoring, and does not equate to theoretical complexity. Nevertheless, it aligns with the design goal that the decoding core scales near-linearly under bounded beam size and candidate limits, and manifests as growth substantially slower than cubic-time methods.

The parameters *α* and *β* control evidence injection and pseudoknot retention, respectively. On the IRIS benchmark, the *α* that optimizes F1, LL, and covariation-based KL varies markedly across transcripts, and these metrics often select different optima, consistent with their targeting distinct signals (LL local accessibility, KL evolutionary constraint, and PARIS/chimeric evidence pairwise interactions). Thus, *α* is best treated as an objective-specific tuning knob chosen to match the application. By contrast, *β* operates only at final selection, biasing retention toward pseudoknotted candidates with no added search cost. On ArchiveII, *β* = 0 yields conservative pseudoknot calling (high precision, low recall), whereas increasing *β* improves pseudoknot-detection F1 with only a modest reduction in overall structure F1, providing a controllable trade-off between global accuracy and pseudoknot sensitivity.

In *in vivo* settings, minimum free energy (MFE) alone is often an insufficient criterion for structural validity(28). Cellular mechanisms such as ATP-dependent remodeling, RBP-mediated interactions, and co-transcriptional kinetics imply that actual structures may deviate from thermodynamic equilibrium(6). Against this background, in the ZIKV case study where no genome-wide ground truth exists, we use COMRADES-derived chimeric consistency 𝒞 (*S*) as the primary comparison signal and treat thermodynamic energy as a reference axis. The results show that objective-driven tuning of *α* by directly maximizing 𝒞 (*S*) enables CPLfold to improve evidence concordance in a reproducible manner, bypassing the expensive randomized constraint sampling of the COMRADES pipeline and making genome-scale folding more practical on standard computational resources.

Practically, these results suggest the following: for sequence-only folding of long RNAs, CPLfold shows good scalability while maintaining high coverage, with lower computational cost in our benchmarks, supporting its use as a default method. When PARIS/icSHAPE evidence is available but reliable ground truth is lacking, *α* can be selected stably using held-out LL and the resulting gains generalize across sequences. For genome-scale scenarios with high-coverage chimeric evidence such as COMRADES, *α* can be tuned directly by optimizing 𝒞 (*S*), turning evidence concordance into a reproducible optimization criterion. If an application prioritizes pseudoknot detection, grid search over *β* under a fixed computational budget can identify a suitable value to increase sensitivity.

Several limitations remain. First, the current two-phase cascade primarily covers single-layer crossing extensions and cannot represent more complex multi-layer pseudoknots. Second, evidence integration relies on additive support terms and does not explicitly model mapping ambiguity, experimental biases, or structural ensembles, potentially making predictions susceptible to noise when evidence is sparse or conflicting. Third, final rescoring depends on existing pseudoknot-capable energy parameterizations, yet kinetics and protein interactions in *in vivo* environments imply that thermodynamic energy can serve only as a limited reference. Future work may extend Phase 2’s crossing space while preserving the bounded candidate pool and scalability, and incorporate uncertainty-aware evidence weighting and multiobjective tuning strategies to further enhance robustness and interpretability under complex *in vivo* data.

## Author contributions statement

**K.W**.: Writing – original draft, visualisation, validation, software, methodology, investigation, formal analysis, data curation, conceptualisation; **G.K**.: Writing – review & editing, supervision, data, conceptualisation.; **S.C**.: Writing – review & editing, supervision, resources, project administration, conceptualisation.

## ACKNOWLEDGEMENTS

This work was supported by the United Kingdom Research and Innovation (grant EP/S02431X/1), UKRI Centre for Doctoral Training in Biomedical AI at the University of Edinburgh, School of Informatics. We appreciate the provision of compute resources through Isambard AI (University of Bristol / UKRI). We thank Dominik Grabarczyk for reviewing the manuscript and providing helpful comments.

## References

1. David H Mathews. Predicting RNA secondary structure by free energy minimization. Theoretical Chemistry Accounts, 116(1):160–168, 2006.

2. Grzegorz Kudla, Andrew W Murray, David Tollervey, and Joshua B Plotkin. Coding-sequence determinants of gene expression in escherichia coli. science, 324(5924):255– 258, 2009.

3. Omer Ziv, Marta M Gabryelska, Aaron TL Lun, Luca FR Gebert, Jessica Sheu-Gruttadauria, Luke W Meredith, Zhong-Yu Liu, Chun Kit Kwok, Cheng-Feng Qin, Ian J MacRae, et al. Comrades determines in vivo RNA structures and interactions. Nature methods, 15(10):785–788, 2018.

4. Phillip J Tomezsko, Vincent DA Corbin, Paromita Gupta, Harish Swaminathan, Margalit Glasgow, Sitara Persad, Matthew D Edwards, Lachlan Mcintosh, Anthony T Papenfuss, Ann Emery, et al. Determination of rna structural diversity and its role in hiv-1 rna splicing. Nature, 582(7812):438–442, 2020.

5. He Zhang, Liang Zhang, Ang Lin, Congcong Xu, Ziyu Li, Kaibo Liu, Boxiang Liu, Xiaopin Ma, Fanfan Zhao, Huiling Jiang, et al. Algorithm for optimized mrna design improves stability and immunogenicity. Nature, 621(7978):396–403, 2023.

6. Robert C Spitale, Ryan A Flynn, Qiangfeng Cliff Zhang, Pete Crisalli, Byron Lee, Jia-Wei Jung, Hannes Y Kuchelmeister, Pedro J Batista, Eduardo A Torre, Eric T Kool, et al. Structural imprints in vivo decode RNA regulatory mechanisms. Nature, 519(7544):486–490, 2015.

7. Nathan A Siegfried, Steven Busan, Gary M Rice, Julie A E Nelson, and Kevin M Weeks. Rna motif discovery by shape and mutational profiling (shape-map). Nature Methods, 11 (9):959–965, 2014.

8. Katherine E Deigan, Tian W Li, David H Mathews, and Kevin M Weeks. Accurate shape-directed RNA secondary structure prediction. Proceedings of the National Academy of Sciences, 106(1):97–102, 2009.

9. Stefan Washietl, Ivo L Hofacker, Peter F Stadler, and Manolis Kellis. Rna folding with soft constraints: reconciliation of probing data and thermodynamic secondary structure prediction. Nucleic Acids Research, 40(10):4261–4272, 2012.

10. Jianyu Zhou, Pan Li, Wanwen Zeng, Wenxiu Ma, Zhipeng Lu, Rui Jiang, Qiangfeng Cliff Zhang, and Tao Jiang. Iris: a method for predicting in vivo RNA secondary structures using paris data. Quantitative Biology, 8(4):369–381, 2020.

11. Grzegorz Kudla, Sander Granneman, Daniela Hahn, Jean D Beggs, and David Tollervey. Cross-linking, ligation, and sequencing of hybrids reveals rna–rna interactions in yeast. Proceedings of the National Academy of Sciences, 108(24):10010–10015, 2011.

12. Zhipeng Lu, Qiangfeng Cliff Zhang, Byron Lee, Ryan A Flynn, Martin A Smith, James T Robinson, Chen Davidovich, Anne R Gooding, Karen J Goodrich, John S Mattick, et al. Rna duplex map in living cells reveals higher-order transcriptome structure. Cell, 165(5):1267–1279, 2016.

13. Elena Rivas and Sean R Eddy. A dynamic programming algorithm for RNA structure prediction including pseudoknots. Journal of molecular biology, 285(5):2053–2068, 1999.

14. Liang Huang, He Zhang, Dezhong Deng, Kai Zhao, Kaibo Liu, David A Hendrix, and David H Mathews. Linearfold: linear-time approximate RNA folding by 5’-to-3’dynamic programming and beam search. Bioinformatics, 35(14):i295–i304, 2019.

15. Mirela S Andronescu, Cristina Pop, and Anne E Condon. Improved free energy parameters for rna pseudoknotted secondary structure prediction. Rna, 16(1):26–42, 2010.

16. Douglas H Turner and David H Mathews. Nndb: the nearest neighbor parameter database for predicting stability of nucleic acid secondary structure. Nucleic acids research, 38 (Suppl_1):D280–D282, 2010.

17. Chuong B Do, Daniel A Woods, and Serafim Batzoglou. Contrafold: RNA secondary structure prediction without physics-based models. Bioinformatics, 22(14):e90–e98, 2006.

18. Michael Zuker. On finding all suboptimal foldings of an RNA molecule. Science, 244(4900):48–52, 1989.

19. Ignacio Tinoco Jr and Carlos Bustamante. How RNA folds. Journal of molecular biology, 293(2):271–281, 1999.

20. Philippe Brion and Eric Westhof. Hierarchy and dynamics of RNA folding. Annual review of biophysics and biomolecular structure, 26(1):113–137, 1997.

21. Song Cao and Shi-Jie Chen. Predicting structures and stabilities for h-type pseudoknots with interhelix loops. RNA, 15(4):696–706, 2009.

22. Jihong Ren, Baharak Rastegari, Anne Condon, and Holger H Hoos. Hotknots: heuristic prediction of RNA secondary structures including pseudoknots. RNA, 11(10):1494–1504, 2005.

23. Anthony J Travis, Jonathan Moody, Aleksandra Helwak, David Tollervey, and Grzegorz Kudla. Hyb: a bioinformatics pipeline for the analysis of clash (crosslinking, ligation and sequencing of hybrids) data. Methods, 65(3):263–273, 2014.

24. Ronny Lorenz, Stephan H Bernhart, Christian Höner zu Siederdissen, Hakim Tafer, Christoph Flamm, Peter F Stadler, and Ivo L Hofacker. Viennarna package 2.0. Algorithms for molecular biology, 6(1):1–14, 2011.

25. Elena Rivas, Jody Clements, and Sean R Eddy. A statistical test for conserved RNA structure shows lack of evidence for structure in lncrnas. Nature methods, 14(1):45–48, 2017.

26. Michael F Sloma and David H Mathews. Exact calculation of loop formation probability identifies folding motifs in RNA secondary structures. RNA, 22(12):1808–1818, 2016.

27. Stanislav Bellaousov and David H Mathews. Probknot: fast prediction of rna secondary structure including pseudoknots. Rna, 16(10):1870–1880, 2010.

28. Silvi Rouskin, Meghan Zubradt, Stefan Washietl, Manolis Kellis, and Jonathan S Weissman. Genome-wide probing of rna structure reveals active unfolding of mrna structures in vivo. Nature, 505(7485):701–705, 2014.

